# Intracellular Amyloid-β in the Normal Rat Brain and Human Subjects

**DOI:** 10.1101/2023.02.15.528693

**Authors:** Asgeir Kobro-Flatmoen, Thea Meier Hormann, Gunnar Gouras

**Author notes:** Correspondence to: Asgeir Kobro-Flatmoen, Full address: Olav Kyrres gate 9, 7030 Trondheim, Norway.

## Abstract

Amyloid-beta (Aβ) is a normal product of neuronal activity, and the two most common variants are 40 or 42 residues long. Of these, the 42 residue-version (Aβ42) is normally less abundant but more prone to self-aggregate, and is thought to cause Alzheimer’s disease (AD). Much knowledge about AD-pathogenesis comes from the study of rodents made to model aspects of the disease by expressing AD-relevant human transgenes, like human amyloid precursor protein (APP) containing mutations that drive up Aβ production or increase the Aβ42/40 ratio and thereby causes AD. Yet, when it comes to the normal expression of Aβ42 in rodent brains, surprisingly little is known. Here we characterize the expression of Aβ42 throughout the brain of normal, outbred Wistar rats, including animals from 3-18 months of age. We find that intracellular Aβ42 (iAβ42) is present in neurons located throughout the brain at all ages of normal Wistar rats, but that the levels vary greatly between brain regions. In cortex, we observe the highest levels of iAβ42 in neurons that are part of layer II of the entorhinal cortex (EC), along with neurons in the hippocampus, closely followed by neurons in the somatosensory cortex. Among subcortical structures, we observe the highest levels iAβ42 in the locus coeruleus. In order to explore whether the striking presence of iAβ42 in rat EC also holds true in human subject free of neurological disease, we examined EC of six such cases ranging from ages 20-88 years. In all six cases, we find that iAβ42 is present in EC layer II-neurons.

Our findings support two conclusions about iAβ42. First, iAβ42 is present in neurons of wild-type Wistar rats and is restricted to the same structures where iAβ accumulates, and Aβ-plaques form, in a much used AD model based on Wistar rats (the McGill-R-Thy1-APP rat model). The difference between wild-type Wistar rats and these AD model rats, with respect to Aβ42, is therefore a quantitative one rather that a qualitative one. This indicates that the McGill rat model in fact models the underlying wild-type neuronal population-specific vulnerability to Aβ42-accumulation. Second, because the McGill rat model closely mimics the human AD-associated spatiotemporal sequence of amyloid plaque deposition, this model may offer a useful representation of the pre-plaque neuronal accumulation of iAβ42. Our findings in human cases are in line with prior findings, and substantiate the notion that neurons in layer II of EC are vulnerable to accumulation of iAβ42.

## Introduction

Amyloid-beta (Aβ) is a small peptide and a normal product of neuronal activity^1-4^. Several variants of Aβ exist, and all result from sequential enzymatic cleavage of the amyloid precursor protein (APP) ^5^. The two most common variants of Aβ are 40 or 42 residues long ^6-9^, and of these two, the 42 residue-version (Aβ42) is normally less abundant but more prone to self-aggregate, owing to its longer C-terminus that makes the peptide more hydrophobic ^10^. Changes in the amount or form of Aβ in the brain, predominantly involving Aβ42, play a decisive role in the development of Alzheimer’s disease (AD) ^11-13^, which is the most common neurodegenerative disease leading to dementia. The onset of AD is characterized by accumulation of intracellular Aβ42 (iAβ42), deposits of extracellular amyloid plaques, hyperphosphorylation of the protein tau (p-tau) that subsequently forms neurofibrillary tangles (NFTs), and severe neuronal death. Three of these pathologies, namely accumulation of iAβ42, formation of NFTs and severe neuronal death, are predominantly restricted to the medial temporal lobes during the initial stages of the disease. Meanwhile, amyloid plaques tend to arise in neocortical areas^14^. Thus, while Aβ likely plays a crucial role in the development of AD, neither its physiological role nor its normal expression in neurons is fully understood^15,16^.

A large body of knowledge about AD-pathogenesis comes from the study of rodents made to model aspects of the disease by expressing AD-relevant human transgenes. For example, use of such rodent models established that early changes involving Aβ42 are followed by changes involving p-tau (reviewed in) ^17^, and that spreading along axonal tracts- and seeding of further pathology is a capacity inherent to both Aβ^18,19^ and p-tau^20,21^. A series of experimental results from Aβ-immunization on a transgenic rodent model forming both Aβ and tau pathology have furthermore shown that, when reducing Aβ42 early on in the pathological cascade, cognitive impairment is reduced or prevented. Once tau-pathology has formed however, reducing Aβ42 is no longer sufficient to prevent cognitive impairment^22,23^. The latter finding is one that still offers hope for early intervention by Aβ-immunization trials.

While much has come from studying rodent models, it is important to keep in mind that these models of AD are not fully representative of the disease. But it is also important to realize that such models likely do offer a unique window into aspects of pathology at prodromal stages of the disease. We and others^17^ therefore believe that rodent models represent opportunities to understand crucial, disease-initiating events, in particular events that involve Aβ42. It follows that in order to understand what early pathological changes may represent, one must understand the normal condition to which the comparison is made. Yet, when it comes to the normal expression of Aβ42 in rodent brains, surprisingly little is known.

A well-established model for the spatiotemporal onset- and progression of amyloid pathology is the Wistar-based, transgenic McGill-R-Thy1-APP rat model, hereafter referred to as the McGill rat model^24-26^. We therefore decided to characterize the expression of Aβ42 throughout the brain of normal, outbred Wistar rats, ranging from the time when they are young adults (three months) up to and including relative old age (18 months). We find that intracellular Aβ42 (iAβ42) is present in neurons located throughout the brain at all ages of normal Wistar rats, but that the levels vary greatly between brain regions. In cortex, we observe the highest levels of iAβ42 in neurons that are part of layer II of the entorhinal cortex (EC), along with neurons in the hippocampus, closely followed by neurons in the somatosensory cortex. Among subcortical structures, we observe the highest levels iAβ42 in the locus coeruleus. In order to explore whether the striking presence of iAβ42 also holds true in cognitively intact humans, we examined EC of six cases ranging from ages 20-88. In all six cases, we find that iAβ42 is present in EC layer II-neurons.

Our findings support two conclusions about iAβ42. First, iAβ42 is present in neurons of wild-type Wistar rats and is restricted to the same structures where iAβ accumulates, and Aβ-plaques form, in the McGill AD rat model. The difference between wild-type Wistar rats and McGill model rats, with respect to Aβ42, is therefore a quantitative one rather that a qualitative one. This indicates that the McGill rat model in fact models the underlying wild-type neuronal population-specific vulnerability to Aβ42-accumulation. Second, because the McGill rat model closely mimics the human AD-associated spatiotemporal sequence of amyloid plaque deposition^26^, this model may offer a useful representation of the pre-plaque neuronal accumulation of iAβ42. Our findings in human cases are in line with prior findings, and substantiate the notion that neurons in layer II of EC are vulnerable to accumulation of iAβ42.

## Methods

For this study we used 12 outbred Wistar Han rats. Nine were bred at the Kavli Institute for Systems Neuroscience, originating from a pair purchased from Charles River (France), while three were purchased from Charles River directly. The animals were divided into four age-groups, including a 3-month (M) group (two males, one female), a 6 M group (three males), a 12 M group (one male, two females), and an 18 M group (three females). All procedures were approved by the Norwegian Animal Research Authority, and follow the European Convention for the Protection of Vertebrate Animals used for Experimental and Other Scientific Purposes. We kept the animals on a 12-h light/dark cycle under standard laboratory conditions (19–22 °C, 50–60%humidity), with access to food and water ad libitum. Use of the archival human formalin-fixed postmortem brain tissues was approved by the Ethical review authority (Etikprövningsmyndighetens), Sweden (permit dnr 2021-01609). The human cases, with pathological remarks, include: Case 1, 20 year old male (no abnormalities mentioned); 2 Case 2, 56 year old female (no significant abnormalities); Case 3, 64 year old male (emphysema); Case 4, 73 year old male (pneumonia); Case 5, 79 year old male (hearth disease); Case 6, 88 year old male (hearth disease).

### Processing and immunohistochemistry

Animals were fully anaesthetised by placement in a chamber containing 5% isoflurane gas (Baxter AS, Oslo, Norway) and then given an intraperitoneal injection of pentobarbital (0.1ml per 100g; SANIVO PHARMA AS, Oslo, Norway). Transcardial perfusions were carried out with room tempered Ringer’s solution (in mM: NaCl, 145; KCl, 3.35; NaHCO3, 2.38; pH ∼6.9) immediately followed by circulation of 4% freshly depolymerised paraformaldehyde (PFA; Merck Life Science AS/Sigma Aldrich Norway AS, Oslo, Norway) in a 125 mM phosphate buffer (PB) solution (VWR International, Pennsylvania, USA), pH 7.4. The brains were extracted and post fixed overnight in PFA at 4°C, and then placed in a freeze protective solution (dimethyl sulfoxide in 125 mM PB with 20% glycerol, Merck Life Science AS/Sigma Aldrich Norway AS, Oslo, Norway) until sectioning. Brains were sectioned coronally at 40 μm in six series, using a freezing sledge microtome.

### Rat tissue

Immunohistochemistry was done on free-floating sections. To enable optimal access to the antigen the tissue was subjected to gentle heat induced antigen retrieval by placement in PB at 60°C for two hours. To prevent unspecific labelling the sections were incubated for 2 × 10 minutes in PB solution containing 3% hydrogen peroxide (Merck Life Science AS/Sigma Aldrich Norway AS, Oslo, Norway), and then incubated for 1 hour in PB solution containing 5% normal goat serum (Abcam Cat# ab7481, RRID: AB_2716553). The sections were then incubated with the primary antibody, IBL rabbit anti-Aβ42 (C-terminal specific, from Immuno-Biological Laboratories Cat# JP18582, RRID: AB_2341375) ^27,28^, at a 1:500 concentration, in PB containing 0.2% saponin (VWR International AS, Bergen, Norway) for approximately 65 hours (three days). Note that we found that incubation for 20 hours (overnight) gave the same qualitative labelling in the sense that we found the same relative difference in signal between distinct neuronal populations, but that the longer incubation-time increased the overall labelling. Next, sections were rinsed for 3 × 5 minutes in Tris-buffered saline (50 mM Tris, 150 mM NaCl, pH 8.0) containing 0.2 % saponin (TBS-Sap), and then incubated with biotinylated secondary antibody goat anti-rabbit (1:400; Sigma-Aldrich Cat#B8895, RRID: AB_258649), in TBS-Sap for 90 minutes at room temperature. Sections were then rinsed in TBS-Sap, incubated in Avidin-Biotin complex (ABC, Vector Laboratories Cat# PK-4000, RRID: AB_2336818) for 90 minutes at room temperature, rinsed again, and then incubated in a solution containing 0.67% 3,3’-Diaminobenzidine (Sigma/Merck, Cat# D5905) and 0.024% hydrogen peroxide for 5 minutes. Sections from animals ranging across the age groups were incubated together in order to safeguard against possible confounds stemming from potentially subtle differences in incubation-time or the quality of the solutions.

### Human tissue

Blocks containing EC were embedded in paraffin, sectioned coronally at 8μm, and mounted on glass slides. Prior to immunohistochemical procedures, the tissue was deparaffinised by immersion in xylene for 5 minutes x 3 times. The tissue was then rehydrated by being dipped 10 times in water containing decreasing amounts of ethanol (100%, 90%, 70%, 50%, 0%). We blocked against unspecific antigens by applying DAKO Real blocking solution (DAKO, Cat# S2023) for 10 minutes, followed by application of normal goat serum (same as above) for 1 hour. To detect Aβ42, we used the same anti-Aβ42 antibody in the same solution as listed above, and incubated the tissue overnight. To also test for the presence or reelin, we used a mouse anti-reelin antibody (Millipore, Cat# MAB5364; RRID:AB_2179313) and carried out double-immunoenzyme labelling. For the anti-Aβ42 antibody we thus used Mach 2 Rabbit HRP Polymer antibody (Biocare Medical, Cat# RHRP520) as our secondary antibody, and visualized it with AEC Red (Sigma-Aldrich, Cat# AEC101). For the anti-reelin antibody, we used a Mach 2 Mouse AP-Polymer (Biocare Medical, Cat# MALP521G) as secondary antibody, and visualized it with Ferangi Blue Chromogen (Biocare Medical, Cat# BRR813AS). Nissl staining was done by dehydrating sections in ethanol, then clearing in xylene, before rehydration in de-ionized water and staining with Cresyl violet (1 g/L) for ∼5 minutes. Sections were then alternately dipped in ethanol–acetic acid (5 mL acetic acid in 1 L 70% ethanol) and rinsed with cold water until we obtained the desired differentiation. The sections were then dehydrated, cleared in xylene and coverslipped with Entellan (Merck KGaA, Darmstadt, Germany).

Processed rat tissue was mounted in Tris-buffered saline on Superfrost® Plus slides (Thermo Scienfific, Menzel Gäser). A subset of sections was counterstained with Cresyl violet (Nissl-stain). Mounted sections were left to dry overnight and then coverslipped (24×60, Thermo Scienfific, Menzel Gäser) using a Xylene- (VWR International AS, Bergen, Norway) and Entallan solution (Merck, Darmstadt, Germany). Processed human tissue was left to dry on a heating plate at 37°C for 30 minutes, and the coverslipped using a water-based medium (Fluoromount-G^®^, Southern Biotech, Cat# 0100-01).

### Imaging

All rat sections were scanned under identical settings using an automated Zeiss Axio Scan.Z1 system, with a 20x objective (Plan-Apochromat 20x/0.8 M27). The analyses were done in Zen software (2.6, Blue Ed.), based on a qualitative assessment of the intensity of the signal. We divide the intensity of the signal, from here on referred to as the ‘level(s)’ of Aβ42, into five categories: none, denoted -; minimal levels, denoted (+); low-to-moderate levels, denoted +; moderate-to-high levels, denoted ++; and high levels, denoted +++. Representative examples for each level are shown in Figure 1. Human tissue was inspected using a Zeiss Axio Imager.M1 microscope, and overview images were acquired with this microscope by a MBF Bioscience CX9000 camera using a 10X objective (Zeiss, N.A. 0.45), while insets were acquired using a 20X objective (Zeiss, N.A. 0.8) and a 100X objective (Zeiss, N.A. 1.4 Oil).

**Figure 1.**
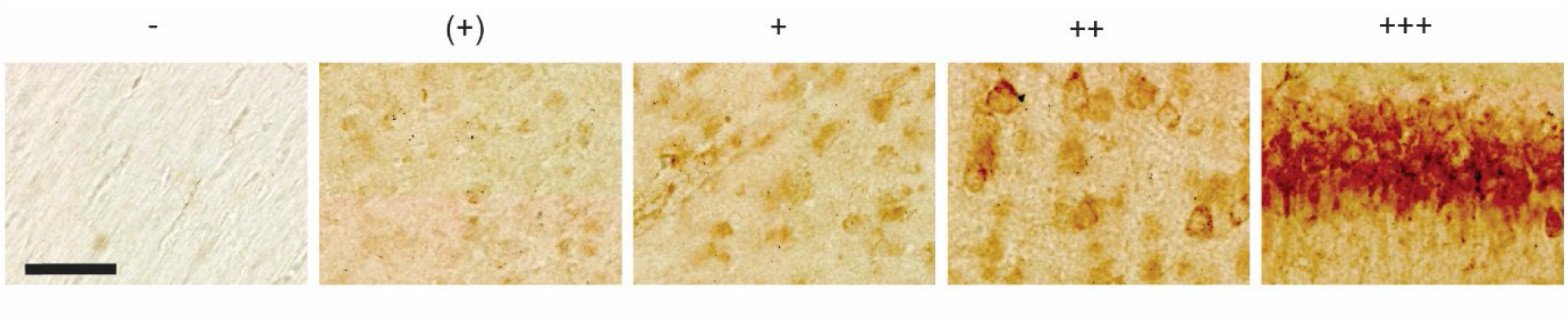
Representative examples of levels of Aβ42 as placed into five different categories, including none (denoted -), minimal levels (denoted (+)), low-to-moderate levels (denoted +), moderate-to-high levels (denoted ++), and high levels (denoted +++). These examples are taken from a 12 month old animal. Scalebar = 50μm.

## Results

To characterize neuronal expression of Aβ42-peptides throughout the brain of normal, outbred Wistar rats, we chose an antibody that has previously been shown to be specific against Aβ42, namely the IBL Aβ42-antibody^28^. For the purpose of validating this antibody also in our hands, we first immunolabeled hippocampal tissue from three-month-old Wistar rats together with hippocampal tissue from six and 12 month old APP KO mice^29^. This experiment demonstrates that while there are high-levels of iAβ42 immunolabeling in neurons of the CA1-subiculum border region of Wistar rats (see more below), there is none or only faint background iAβ42 immunolabeling in corresponding neurons (or any other neurons) of APP KO mice (Fig. 2; note that there are no APP KO rats available), thus substantiating the specificity of the antibody.

**Figure 2.**
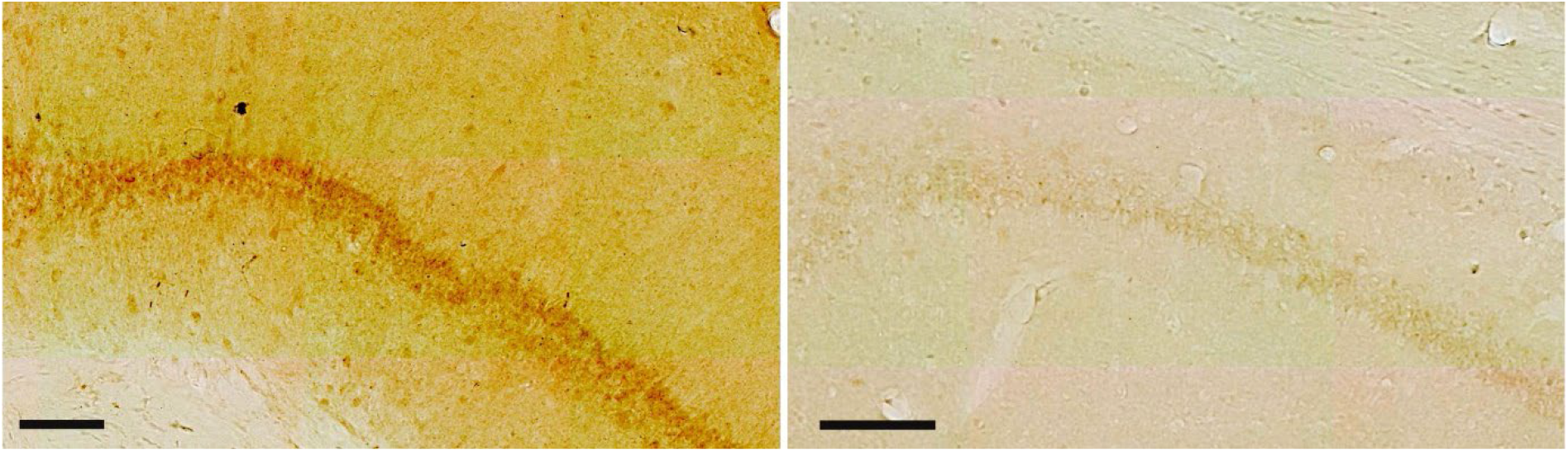
The Aβ42-antibody is specific. (left image) Immunolabeling with the IBL Aβ42-antibody reveals a high level of signal in CA1 of three-month-old Wistar rats, while (right image) practically no signal is present in CA1 of six-month-old Amyloid Precursor Protein-Knockout mice (same results were obtained for 12-month-old Amyloid Precursor Protein-Knockout mice). Scalebars = 100μm. Images were acquired using identical settings.

The following descriptions of our results relate to neuronal somata, unless otherwise specified. In broad terms, intracellular Aβ42 (iAβ42) is present in neurons found throughout the brain at all ages of normal Wistar rats, but the levels vary greatly between regions, and also within certain regions. We find that as a general tendency, iAβ42-levels increase slightly in wild-type Wistar rats up to and including 12 months of age, before dropping slightly from 12 to 18 months. In the forebrain, we observe the highest levels of iAβ42 in parts of the hippocampus and parts of the entorhinal cortex, followed by the primary somatosensory cortex. In the mid- and hindbrain, we observe the highest levels iAβ42 in the locus coeruleus and in Purkinje neurons of the cerebellum.

### Forebrain

#### Hippocampal formation (Archicortex)

Two gradients of iAβ42 stand out in association with the hippocampal formation. One relates to the transverse axis and involves CA1 and the subiculum, while the other relates to the septotemporal axis and involves the whole hippocampal formation. Relating to the transverse axis, high-levels of iAβ42 are present in neuronal somata of the pyramidal layer at the border region between CA1 and the subiculum (CA1/Sub), but the levels of iAβ42 drop when moving away from the CA1/Sub border region. Specifically, in the subiculum, iAβ42-levels are high at the CA1/Sub border region, but this changes to a low-to-intermediate level almost immediately when moving away (distally) from the border region, and remains at this low-to-intermediate level for the remaining distal extent of the subiculum. For CA1, the high iAβ42-levels at the CA1/Sub border region extend into approximately the distal half of CA1. From this point, iAβ42-levels gradually drop as one moves successively further away from the CA1/Sub border region, such that at the proximal extreme of CA1, iAβ42-levels are low (Fig. 3 A). Meanwhile, the gradient relating to the septotemporal axis, which holds true for all the hippocampal fields (DG, CA1-CA3, and Sub), is such that each field has higher levels of iAβ42 in their septal domain *relative to* their temporal domain. While both the transverse and the septotemporal gradient is consistently present across animals, the absolute levels of iAβ42 vary between animals. This between-animal variation is most notable for DG and CA3. Specifically, in some animals the iAβ42-levels in DG and CA3 come close to, though not quite reaching that seen for the CA1/Sub border region, while for other animals the iAβ42-levels in DG and CA3 are substantially lower than that for the CA1/Sub border region (Fig. 3 B).

**Figure 3.**
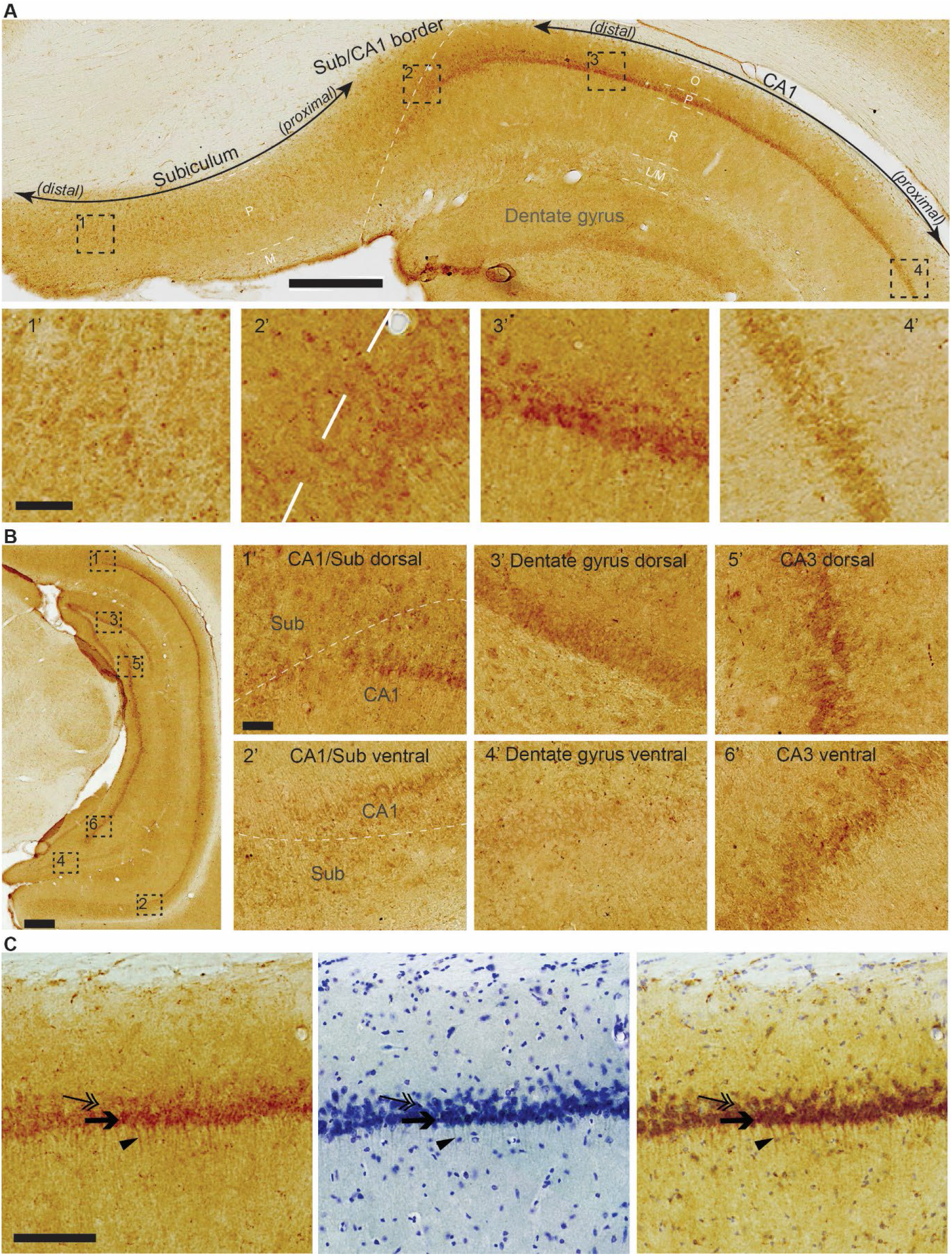
**(A)** Top micrograph: High levels of iAβ42 are present in neuronal somata of the pyramidal layer at the border region between CA1 and the subiculum (distal CA1, proximal subiculum), but iAβ42-levels drop off when moving away from this border region, such that low levels of iAβ42 are present in proximal CA1 and distal subiculum. For the subiculum, the drop in iAβ42-levels occurs almost immediately distal to the CA1/subiculum border region, while for CA1, the drop occurs more gradually when moving increasingly proximal from this border region. Dashed white line indicates border between CA1 and the subiculum. Layers are indicated by short dashed white line and white text for CA1 and Sub: O = Oriens, P = Pyramidale, R = Radiatum, L/M = Lacunosum/Moleculare, M = Moleculare. Bottom micrographs: Higher-powered insets taken from top micrograph, numbers according to location. **(B)** Leftmost micrograph: Hippocampal formation at the maximal dorsoventral extent achieved by a coronal section. Right-side insets: iAβ42-levels are substantially higher in the dorsal than in the ventral portion of the hippocampal formation. Notice the moderate-to-high levels of iAβ42 in dorsal dentate gyrus and dorsal CA3 in this example; this feature is variable between animals. Insets numbered according to location. Dashed white line in 1’ and 2’ indicates the border between CA1 and the subiculum. **(C)** Rightmost micrograph: In CA1, superficially located pyramidal neurons have higher levels of iAβ42 (arrow) than those located deeper (double-arrow), and many of those superficially located CA1 pyramidal neurons have iAβ42 extending into their basal dendrites (arrowhead). Middle micrograph: Nissl stain of adjacent section (Cresyl Violet). Rightmost micrograph: iAβ42-and Nissl stained sections-overlay. Data are from an 18-month-old animal. Scalebars, (A and B) top micrograph 500μm, bottom micrographs, 50μm, (C) 200μm.

The most clearly observable iAβ42-signal is generally restricted to somata with our immunohistochemical method, but CA1 pyramidal neurons constitute an exception in that many show iAβ42 also in their basal dendrites, extending into stratum radiatum. Another feature of CA1 is that iAβ42-levels are noticeably higher in neurons situated in the superficial part of the pyramidal layer; most of the dendritic iAβ42 seems to be associated with these superficially situated neurons (Fig. 3 C).

#### Piriform cortex (Archicortex)

In the piriform cortex, a far greater number of neurons in layer II stain positive for iAβ42 than neurons in layer III. Most neurons have moderate iAβ42-levels, though a few neurons have moderate-to-high levels. Neuropil-labelling is generally low, except in the outer half of layer I, where a moderate Aβ42 positive band of labelling is present (Fig. 4 A, B).

**Figure 4.**
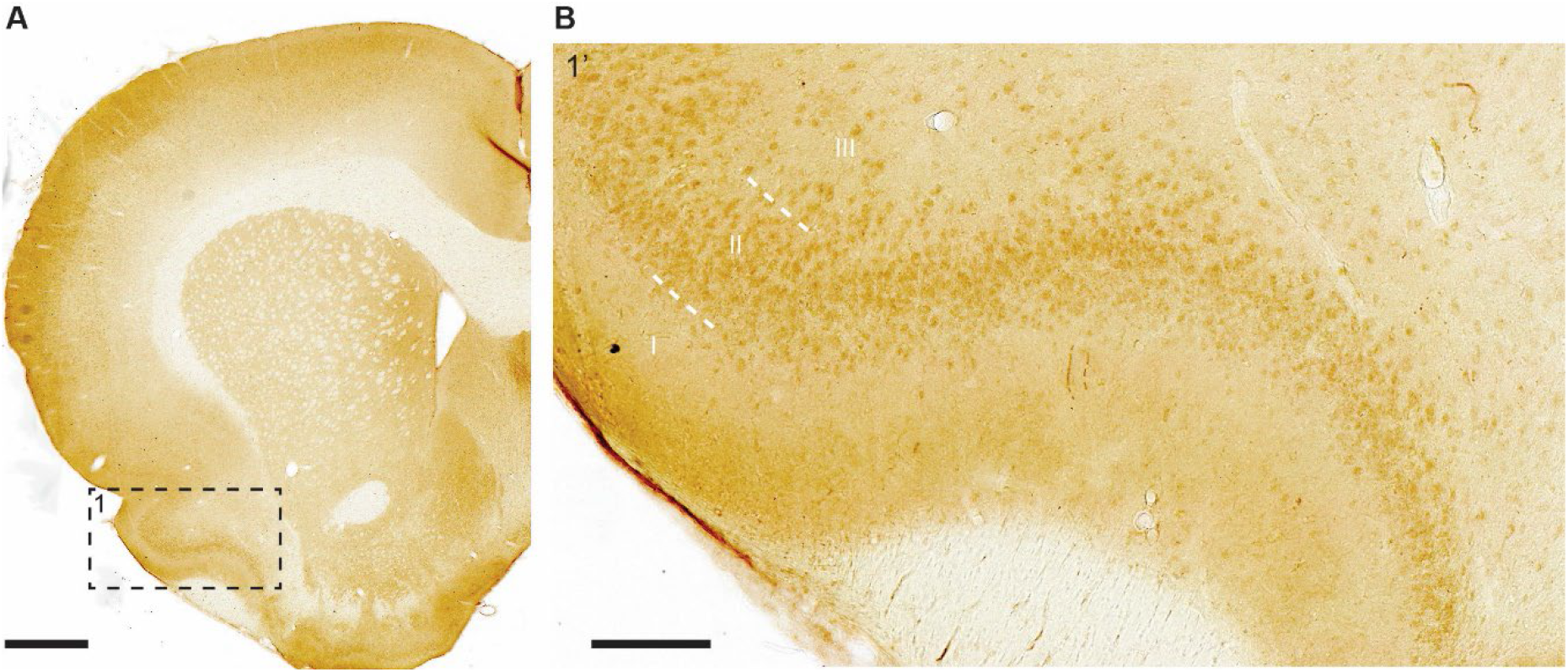
Moderate-to-high levels iAβ42, mainly leaning towards moderate, are present in layer II-neurons of the piriform cortex. A far lesser number of layer III-neurons also stain positive for iAβ42. **(A)** Frontal section with example of piriform cortex indicated (1, dashed rectangle), along with **(B)** higher-powered inset. Note that aside from the signal in neurons, an Aβ42 positive band of labelling is present in the outer half of layer I. Scalebars (A) 1000μm, (B) 200μm.

#### Parahippocampal region (Periallocortex)

In the lateral entorhinal cortex (LEC), the population of large fan neurons that are situated in the superficial part of layer II and close to the rhinal fissure, is one of the few neuronal populations whose somata contain high levels of iAβ42 in the normal rat brain. When moving successively further away from the rhinal fissure, the levels of iAβ42 in fan neurons gradually drop, reaching the lowest levels at the point furthest away from the rhinal fissure (Fig. 5 A). This is different for the medial entorhinal cortex (MEC), where LII contains so-called stellate neurons, which are considered a counterpart population of neurons to the LEC fan-neurons (for more details on properties and differences between LII-neurons in LEC vs MEC, please see the following papers^30,31^). Thus, while a similar gradient to that described for LEC is present also in MEC, with higher levels of iAβ42 being present in stellate neurons close to the rhinal fissure, the total levels of iAβ42 in MEC stellate neurons is generally far less than for LEC fan neurons (Fig. 5 B). However, we observe a notable between-animal variation with respect to iAβ42 in MEC, with occasional animals having moderate-to-high levels in stellate neurons close to the rhinal fissure. In LEC LIII, and to a lesser degree in MEC LIII, neurons close to the rhinal fissure have moderate-to-high levels of iAβ42. Though less pronounced than that for LII, there is a similar gradient vis-a-vis the rhinal fissure, such that neurons located increasingly further away from the rhinal fissure express less iAβ42. We do not observe such a gradient for neuronal somata in deeper layers, where the levels of iAβ42 range from low-to-moderate (Fig. 5 A, B).

**Figure 5.**
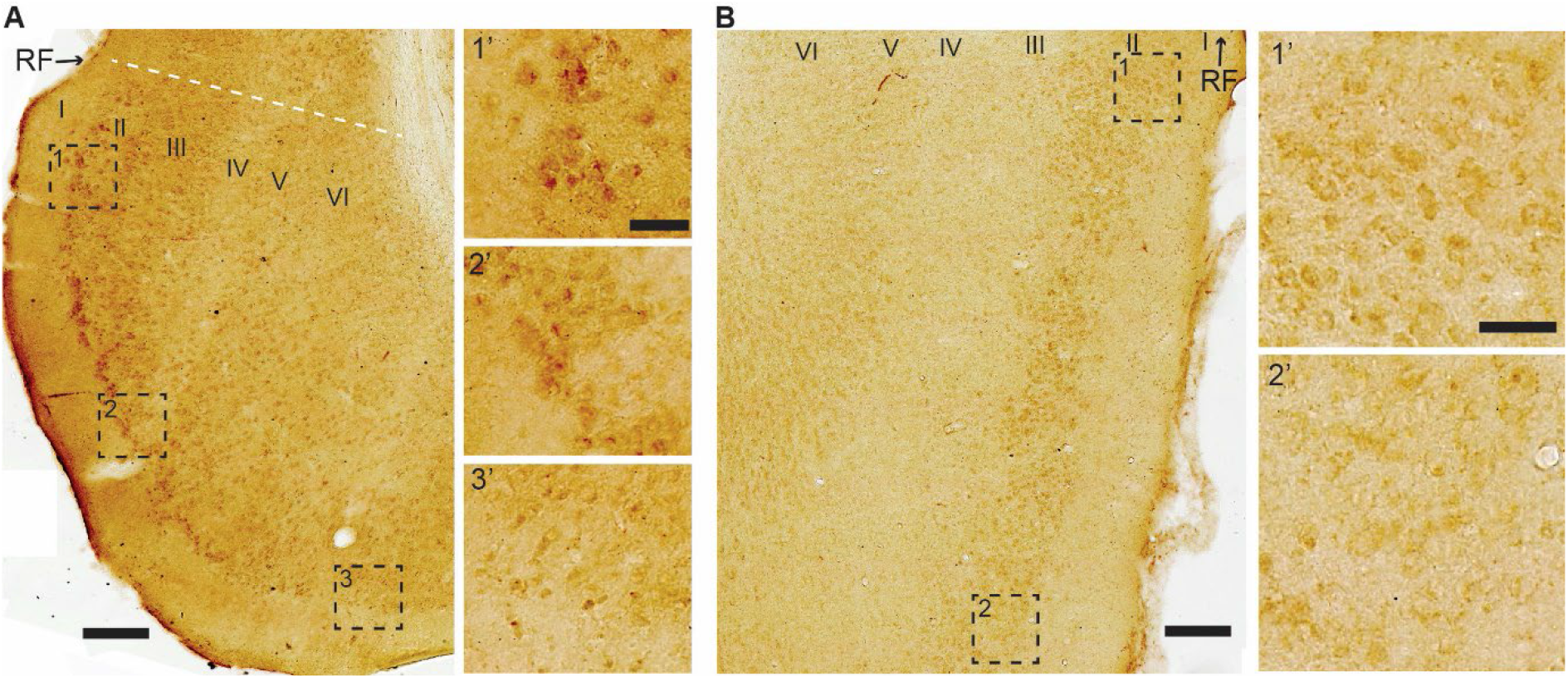
High levels of iAβ42 are present in entorhinal cortex (EC) layer II (LII)-neurons that situate close to the rhinal fissure. **(A)** In lateral entorhinal cortex (LEC), high-levels of iAβ42 are present in neurons in the outer portion of LII that situate close to the rhinal fissure (RF); iAβ42-levels gradually drop when moving successively further away from RF, being reduced to low levels at the point furthest away from RF. This feature is similar, though less pronounced, also in the case of LIII, but is not present in deeper layers. **(B)** In Medial entorhinal cortex (MEC), low-to-moderate levels of iAβ42 is present in LII-neurons located close to RF, and, similar to that of LEC, iAβ42-levels drop when moving successively further away from the rhinal fissure. Layers are indicated (I-VI). Dashed white line in (A) indicates border with perirhinal cortex. Micrographs are from an 18-month-old animal. Scalebars 200μm (insets 50μm).

For the remaining structures that make up the parahippocampal region, namely the pre- and parasubiculum, and the peri- and postrhinal cortices, we find that iAβ42-levels range from low-to-moderate. In the pre- and parasubiculum, low levels tend to dominate in deeper layers while moderate levels tend to dominate in superficial layers. In the perirhinal cortex, the ventrally situated area 35 tends to have low levels of iAβ42 across the layers, while the dorsally situated area 36 tends to have low levels in the superficial layers and low-to-moderate levels in the deeper layers. In the postrhinal cortex, iAβ42-levels are low-to-moderate, with layers II and V-VI tending towards moderate levels, and layer III tending towards low levels.

#### Neocortex (Allocortex)

A majority of neurons in most areas of the neocortex have low or minimal levels of iAβ42. An exception to this general finding is the primary somatosensory cortex, and to a lesser extent the primary motor cortex. The primary somatosensory cortex harbours a great number of neurons with moderate-to-high levels of iAβ42 across layers II-VI, of which those with the highest levels constitute pyramidal neurons in layer V. In the adjacent primary motor cortex, low-to-moderate iAβ42-levels are present in layers II-V, while low levels are present in layer VI (Fig. 6 A-C). Low or low-to-moderate levels of iAβ42 are present also in many layer V pyramidal neurons of the secondary motor cortex and the parietal association cortex; in both of the latter cortices, a lesser number of layer V pyramidal neurons have moderate-to-high iAβ42-levels.

**Figure 6.**
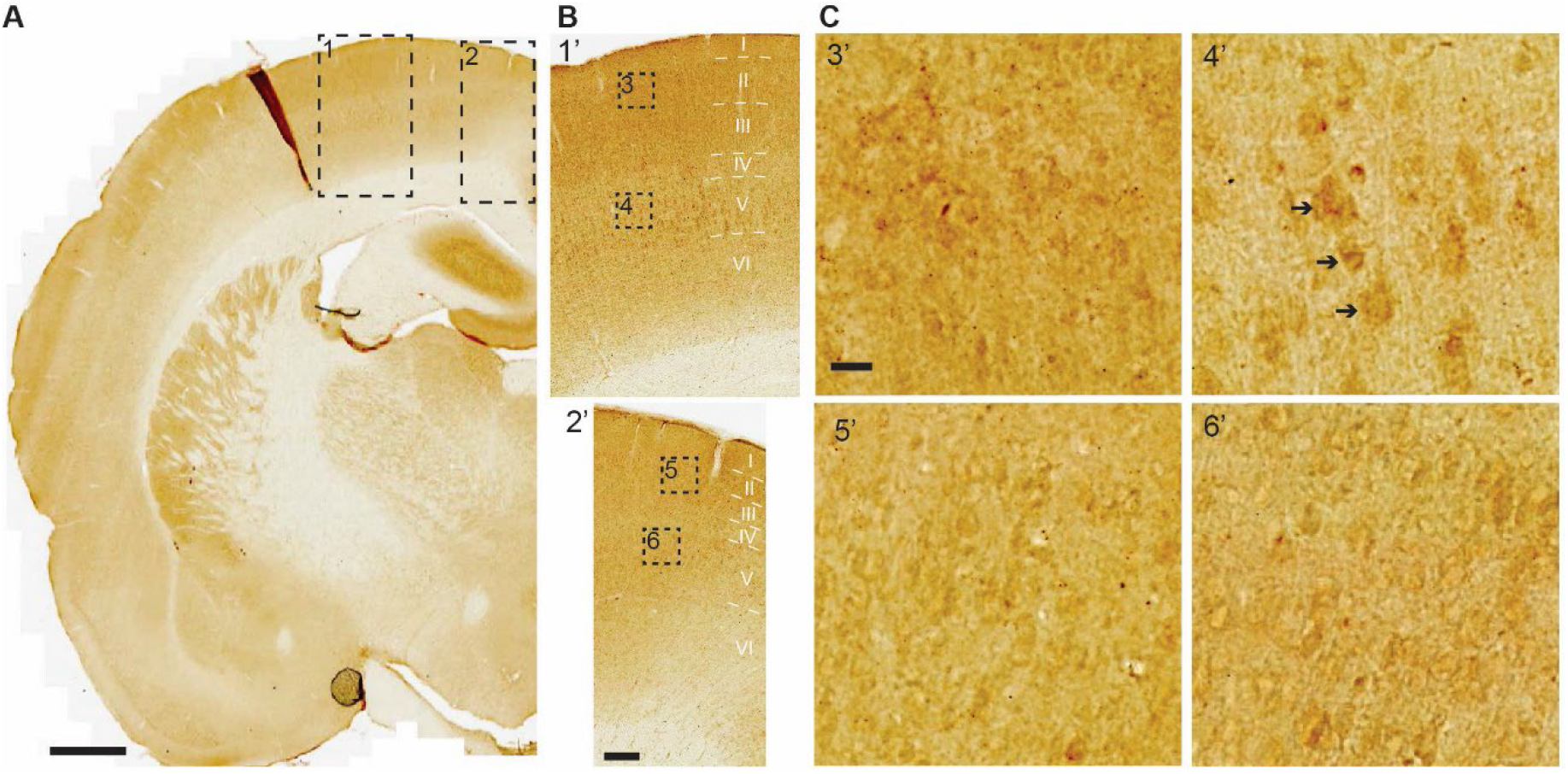
Moderate-to-high levels iAβ42 are present in the primary somatosensory cortex, most notably in layer V. **(A)** Levels of iAβ42 are generally low in across the neocortex. But in the primary somatosensory cortex (dashed rectangle 1), moderate-to-high iAβ42-levels are present in layer V pyramidal neurons, and low-to-moderate levels of iAβ42 are present in neurons of superficial layers along with layer VI. In the adjacent primary motor cortex (dashed rectangle 2), low-to-moderate iAβ42-levels are present in layers II-V, while low levels are present in layer VI. **(B)** and **(C)** shows increasingly higher-powered insets; note the examples of clearly stained layer V pyramidal neurons (arrows) in primary somatosensory cortex (inset 4’). Scalebars (A) 1000μm, (B) 200μm, (C) 20μm.

### Mid- and Hindbrain

In the mid- and hindbrain, levels of iAβ42 are generally minimal. However, striking exceptions to this general finding include the cerebellum, and the locus coeruleus of the brainstem.

### Cerebellum

Most neurons in the cerebellum have no detectable or minimal levels of iAβ42. Purkinje neurons constitute an exception as these have high-levels of iAβ42 and thus stand out in striking contrast to the cerebellar granule neurons (Fig. 7). We do not observe any clear gradient with respect to the distribution of iAβ42-levels between Purkinje neurons of different portions in the cerebellum.

**Figure 7.**
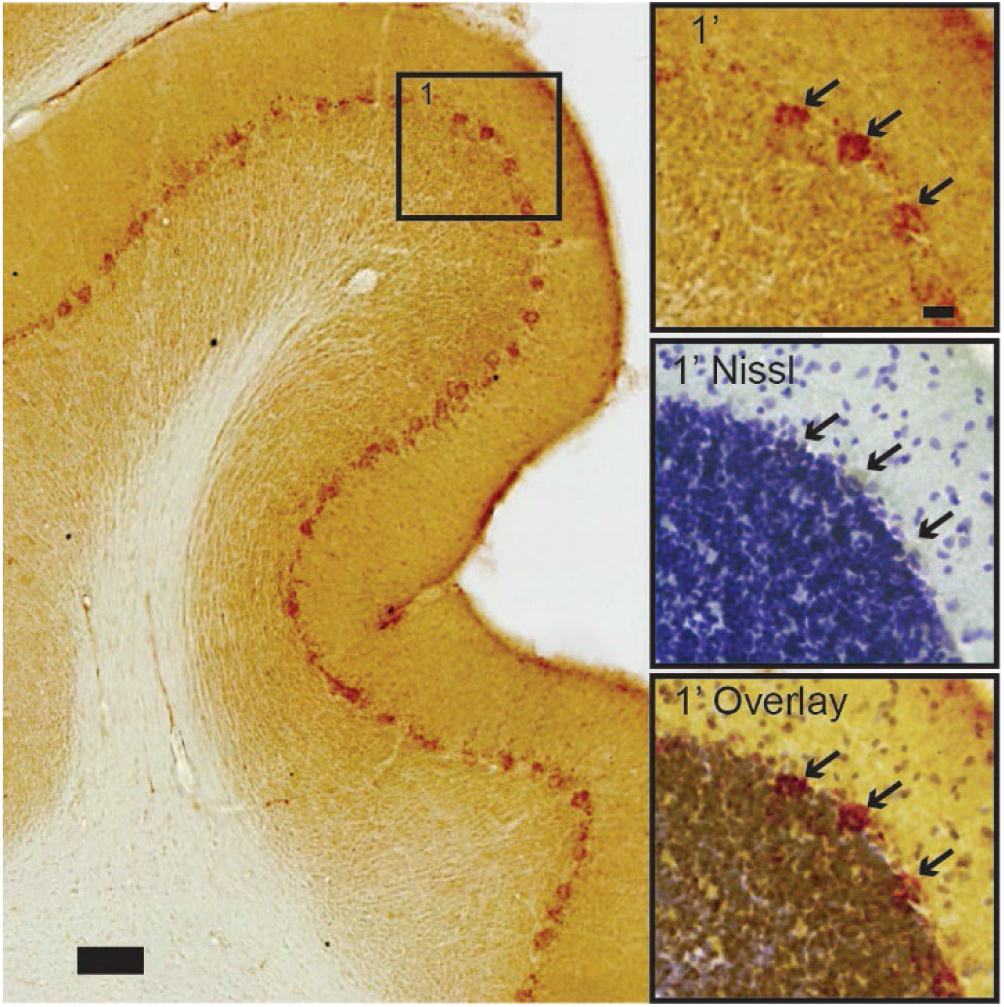
Cerebellar Purkinje neurons contain high-levels of iAβ42, in stark contrast to surrounding neurons. Insets show arrows pointing to (top) iAβ42-positive Purkinje neurons, followed by (middle) the position of these neurons after Nissl counterstain on the same section, and (bottom) a contrast enhanced overlay of the top and middle insets. Scalebars 100μm (top inset 20μm, same for all).

### Locus Coeruleus

In the locus coeruleus, we find high-levels of iAβ42 in neurons located in the very densely packed dorsomedial portion (Fig. 8). A Nissl counterstain reveals that these neurons are round or ovoid, which corresponds to previously published findings about the morphology of neurons in this portion of the locus coeruleus^32^.

**Figure 8.**
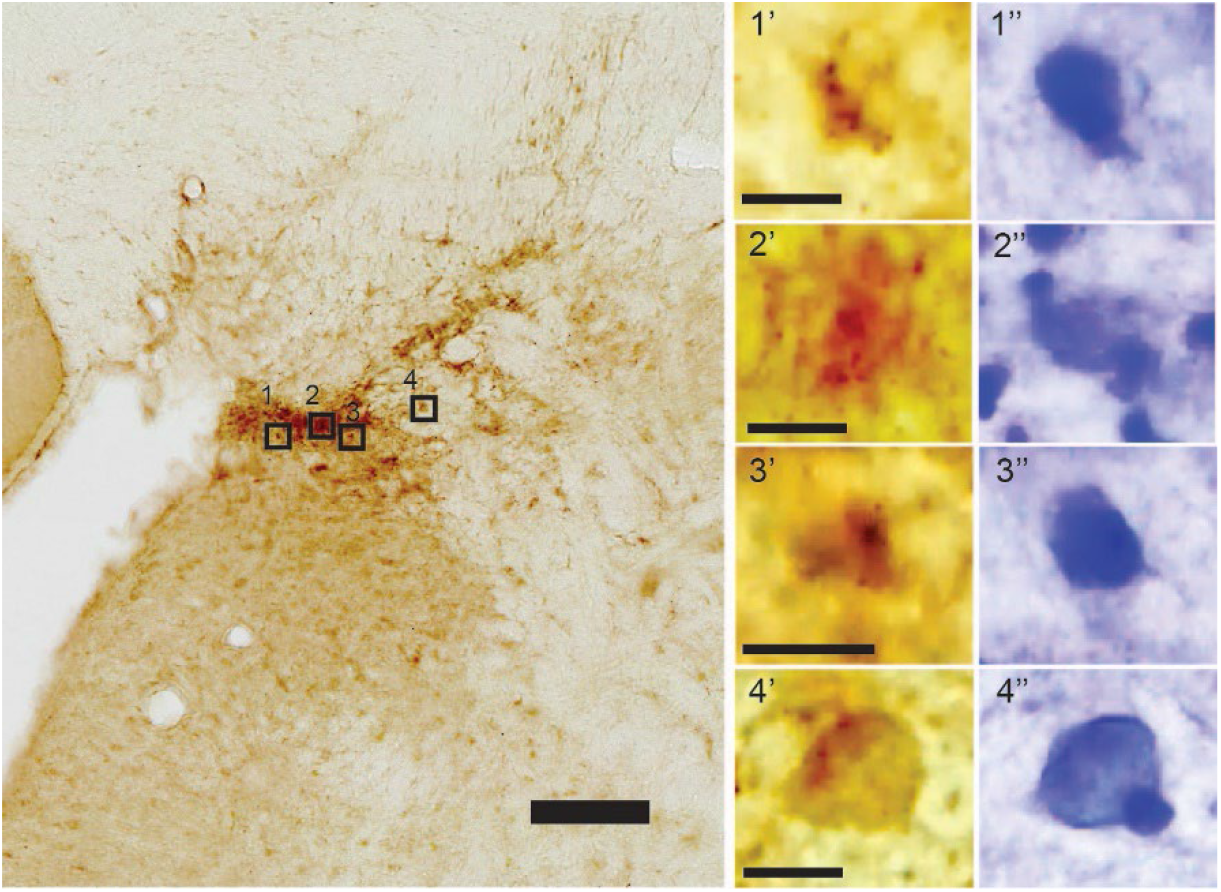
High levels of iAβ42 in cells of the dorsomedial part of the locus coeruleus. A Nissl counterstain done on top of the DAB-labeled section (insets) reveals that several of these cells are round or ovoid. Note that at this portion of the locus coeruleus the cells are very densely packed, and contrast enhancement was necessary to separate adjacent cells. Scalebars 100μm (insets 10μm).

### Human entorhinal cortex

Based on our findings in normal rats, together with multiple lines of evidence showing that accumulation of iAβ42, formation of NFTs and severe neuronal death arises in EC-layer II already at the initial stages of AD, we sought to examine sections of EC from human subjects free of neurological disease. We examined six such cases, ranging from 20-88 years of age. From each human case we immunolabeled sections of EC using the same iAβ42-antibody as that for rats. Similar to our findings in the normal rat brain, we find that iAβ42 is present in EC layer II-neurons in all six human control cases (Fig. 9 A-C). Furthermore, the intensity of iAβ42-labeling was greater in older subjects relative to younger ones. In order to answer whether Aβ42 was located inside EC layer II-neurons also in the oldest case, where some of the Aβ42-labeling started to take on the appearance of small plaques, we counterstained a section with cresyl violet. This revealed that, in many cases, a nucleus and a nucleolus was still situated inside the Aβ42-labeling (Fig. 9 C).

**Figure 9.**
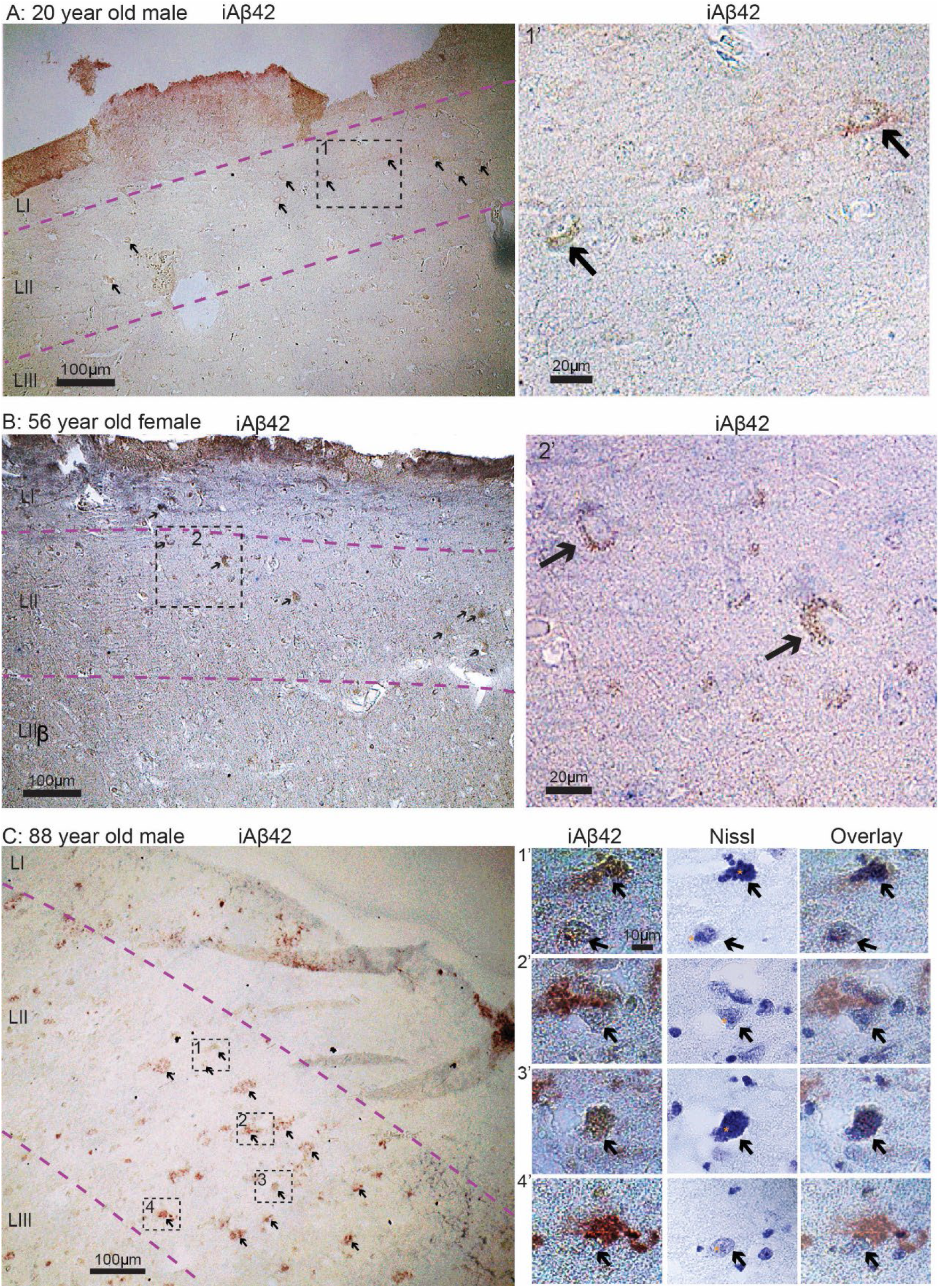
iAβ42 is present in entorhinal cortex layer II-neurons in human subjects free of neurological disease. **(A)** 20-year-old male. **(B)** 56-year-old female. **(C)** 88-year-old male. Overview images in (A-C) were taken with a 10x objective, insets for (A) and (B) were taken with a 20x objective, while insets for (C) were taken with a 100x Oil immersion objective. Immunolabel and Nissl staining is indicated for each image/panel. Stippled lines in light magenta indicate layers. To indicate nucleoli in the middle panels of (C), we have place a small asterisk immediately to the left of each. Scalebars are indicated for each image/panel (in (C), the scalebar in top left image applies to all in this panel).

## Discussion

A generic criticism against transgenic rodent models for AD is that they typically have “non-physiological expression” of their transgenic product or that they fail to form pathology in the correct brain structure^33^. Given that they constitute the vast majority of AD models, it is certainly relevant to address such criticism with respect to rodent models expressing mutant human APP to cause increased levels of Aβ (APP-models). While with the development of AD, Aβ-levels are no longer physiological, which is what rodent APP-models model, it is crucial to know whether a given transgene expresses in the relevant brain structures, but also whether the given transgene causes increased expression of its protein product in structures that ought not to be affected. Yet, little is known about the normal, brain structure-specific expression of Aβ in rodents. Here we characterized the brain-wide presence of iAβ42 in normal outbred Wistar rats. This rat-strain forms the background-strain for the much used McGill rat model, an APP-model that mimics the spatiotemporal sequence of amyloid plaque deposition seen in human AD-subjects^26^ and has an extended pre-plaque period of iAβ-accumulation^25^.

We find that in normal wild-type Wistar rats, neurons in a subset of structures involving both the forebrain and the hindbrain express surprisingly high levels of iAβ42. These structures include the part of LEC LII located close to the rhinal fissure, the CA1/Sub border region, and the somatosensory cortex (forebrain), along with the cerebellum and the locus coeruleus (hindbrain). With respect to the forebrain, these relatively high iAβ42-expressing structures are the exact structures in which amyloid plaque-pathology first arise in the transgenic mutated hAPP-based McGill rat model^26^. In particular, plaque-pathology in the McGill rat model begins in the proximal portion of the subiculum and typically also involves the distal part of CA1 (*see Fig. 12 L&M in*^*26*^), thus perfectly matching the portion of the hippocampus where, relative to other areas, high levels of endogenous iAβ42 are present in normal Wistar rats. Subsequent to the CA1/Sub, plaques in the McGill model deposit in the part of LEC located close to the rhinal fissure (*see Fig. 14 O and 16 O in*^*26*^), again matching the substructure where we find a particularly high-level of endogenous iAβ42 in normal Wistar rats. The same is the case for the neocortex, as the McGill model deposits its first *neocortical* plaques in the somatosensory- and motor cortex, which is the only part of the neocortex in which we find a high level of endogenous neuronal iAβ42 in normal Wistar rats.

We are not aware of any available data regarding the hindbrain of the McGill rat model. Nevertheless, our findings that cerebellar Purkinje neurons and neurons in the locus coeruleus in normal Wistar rats selectively express high levels of iAβ42 are of considerable interest. It is well known that amyloid plaques eventually form in the cerebellum in AD^34^, and such plaques have been linked specifically to Aβ-laden Purkinje neurons^35^. It is notable in this context that even though neurofibrillary tangles do not seem to readily form in the cerebellum, this might depend on the Aβ42-load, since at least one type of familiar AD (PS1-E280A), caused by increased production of Aβ42, leads to formation of cerebellar p-tau^36^. Contrary to this, the locus coeruleus is, alongside laterally situated entorhinal layer II-neurons, particularly prone to early formation of p-tau and formation of neurofibrillary tangles^37^. While little is known regarding the possibility of Aβ accumulating in the locus coeruleus during the pre-clinical phase of AD, a recent study using an antibody against oligomeric or fibrillar Aβ on cases with full-blown AD did report the presence of plaques in the locus coeruleus^38^.

To provide some human correlate to our findings in normal Wistar rats to human brains we also carried out Aβ42-immunolabeling on EC-tissue from six cases of cognitively normal human subjects, ranging from 20-88 years of age. In all six cases, we find iAβ42 in EC layer II-neurons, substantiating prior findings in humans about the vulnerability of such neurons to accumulate Aβ^2,25,39,40^. Why the somata of such EC layer II-neurons (or EC layer II in itself) are not associated with plaque formation in AD remains unclear, although our working hypothesis is that the iAβ42 in somata of EC layer II neurons then shifts to their axon terminals in the perforant pathway which develop prominent Aβ42 aggregation (Roos et al., 2021).

Because the structures with the most notable expression of neuronal iAβ42 in normal Wistar rats are the same as those which show early neuronal iAβ42 in the McGill rat model, and because this rat model closely mimics the AD-related spatiotemporal cascade of AD neuron vulnerability and Aβ-plaque deposition^26^, we conclude that it is well suited for the purpose of modelling the cascade of amyloid pathology seen in the context of AD. We emphasize that this likely includes the pre-plaque neuronal accumulation of iAβ42, which in the McGill rat model extends over approximately 8 months.

## Abbreviations

Aβ: Amyloid-β
AD: Alzheimer’s disease
APP: Amyloid precursor protein
CA1-3: Cornu Ammonis 1-3
EC: Entorhinal cortex
hAPP: human Amyloid precursor protein
iAβ: intracellular Amyloid-β
KO: Knock-out
LII: layer II
LC: Locus coeruleus
LEC: Lateral entorhinal cortex
MEC: Medial entorhinal cortex
NTF: Neurofibrillary tangle
p-tau: hyperphosphorylated tau
Sub: Subiculum

